# A direct computational assessment of vinculin-actin unbinding kinetics reveals catch bonding behavior

**DOI:** 10.1101/2024.10.10.617580

**Authors:** Willmor J. Peña Ccoa, Fatemah Mukadum, Aubin Ramon, Guillaume Stirnemann, Glen M. Hocky

## Abstract

Vinculin forms a catch bond with the cytoskeletal polymer actin, displaying an increased bond lifetime upon force application. Notably, this behavior depends on the direction of the applied force, which has significant implications for cellular mechanotransduction. In this study, we present a comprehensive molecular dynamics simulation study, employing enhanced sampling techniques to investigate the thermodynamic, kinetic, and mechanistic aspects of this phenomenon at physiologically relevant forces. We dissect a catch bond mechanism in which force shifts vinculin between either a weakly- or strongly-bound state. Our results demonstrate that models for these states have unbinding times consistent with those from single-molecule studies, and suggest that both have some intrinsic catch bonding behavior. We provide atomistic insight into this behavior, and show how a directional pulling force can promote the strong or weak state. Crucially, our strategy can be extended to capture the difficult-to-capture effects of small mechanical forces on biomolecular systems in general, and those involved in mechanotransduction more specifically.

## 1 Introduction

Many macromolecules in cells exist under constant bombardment of rapidly changing mechanical forces deriving from their environment [1–3]. These external forces can arise in many ways, for example due to the action of cytoskeletal motors or expansion or compression of biological membranes [1, 2]. While the sources of mechanical force are stochastic, in many cases we can consider the time averaged effect of many such events as creating a static mechanical force on a protein of interest, and ask what is the effect of this constant extensional or compressive force on our molecule of interest [3]. The scale of mechanical forces in biology are minute from a macroscopic perspective, falling in the range of less than one to tens of piconewtons (pN) [4]. While this force magnitude is sufficiently small that we do not expect it to destabilize the folded state of standard proteins [5], specialized proteins called mechanosensors have evolved such that sustained forces of this magnitude (perhaps in a specific direction) can substantially shift their conformational ensemble, resulting in a change of biological function [1–3].

The actin cytoskeleton is a prototypical example of a mechanosensing system due to its role in active mechanical properties of cells including regulating cell shape, motility, and division [6]. Actin filaments are non-covalent semi-flexible polar polymers formed from the end-end arrangement of globular actin monomers [6, 7]. Actin monomers and filaments have dozens of known binding proteins (ABPs) that control an entire dynamical ecosystem through either (i) regulation of the assembly and disassembly of individual filaments,(ii) crosslinking of filaments into networks, (iii) or actively exerting pulling forces on filaments [7, 8]. There are now several examples of how forces on ABPs or on actin filaments can change binding affinity of an ABP to filaments, resulting in mechanical feedback loops that help quickly toggle the local morphology of the network [9–12].

When a force is applied across the junction of actin with an ABP, we expect that interaction to be destabilized. Naively, we would assume that pulling on a protein-protein interaction tilts the energy landscape towards the unbound state, and lowering the barrier towards unbinding [3]; in this scenario simple approximations lead to the “Bell’s law” [13] prediction that pulling with force *F* results in an exponential decrease in the lifetime of the association *τ*_off_,

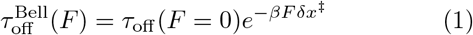

where δ*x*^‡^ represents the “distance to the transition state”.

In reality, the situation is much more complex, since we must consider a free energy surface (FES) for the unbinding reaction, which takes into account a Boltzmann weighted average over all possible configurations of the system. The FES 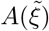 along a reaction coordinate 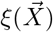 (e.g. number of contacts between a ligand and a protein) is computed through taking a Boltzmann weighted average over all possible configurations. If a pulling force *F* applied along the coordinate *Q* (e.g. distance of the ligand to the center of its binding pocket) is included, the FES as a function of force,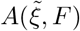 can be expressed as

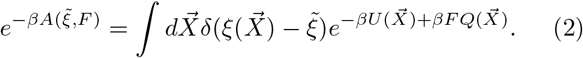

Here, *β* = 1*/*(*k*_B_*T*), *k*_B_ is Boltzmann’s constant, *T* is the temperature, δ is the Dirac delta function, *U* is the potential energy function for the system, and 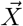 represents the configuration of the system in all 3*N* Cartesian dimensions.

The effect of force on the FES can be complex, because many configurations map to a single value of 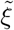, while these can each differ in their value of the pulling coordinate *Q*. This complex dependence is sufficient to explain catch bonding behavior, which would occur when the effective free energy barrier to unbinding increases for larger force. Starting in the early 2000s, with the advance of experimental techniques able to probe unbinding processes with high precision (mainly through microfluidic flow chambers or single molecule force spectroscopy apparatuses), this phenomenon has been found in an increasing number of systems involved in adhesion [14, 15], blood coagulation [16], and mechanotransduction [15, 17, 18].

A simple biochemical framework to explain catch bonding is that of allostery. In this framework, one part of a protein acts to make the binding of the whole protein to its partner constitutively weak. When force is applied to this allosteric modulation domain in the correct direction, the inhibition is released and the protein can reach a higher affinity state with a corresponding longer lifetime. This can be captured using a three-state model where the force activated state has a larger barrier to unbinding than does the zero-force state [19–24].

One example system where catch bonding behavior is believed to follow from this allosteric mechanism is the interaction of the protein vinculin with actin filaments [17], which evolved to promote connection between adhesions and actin filaments that are flowing backwards near a lamellipodium, where the polymerization of actin is used to push forward the cellular membrane, promoting formation of focal adhesions [17, 25]. It is important that this stronger adhesion remains transient, otherwise it would be difficult for the network to dynamically reconfigure itself [25]. The actin binding domains of related proteins talin [26] and *α*-catenin [18] also exhibit catch bonding behavior, and the proposed allosteric mechanism in that case involves transition from a five-helix weak binding state to a four helix strongly bound state, where one part of the bundle has detached and become disordered.

This mechanism is indirectly inferred from experimental measurements, because it has been nearly impossible to directly isolate structures (e.g. when using X-ray crystallography or electron cryomicroscopy (CryoEM)) that are only activated upon applied force, although emerging approaches may enable this going forward [27]. One can try to mimic the allosteric effect of force using mutations on the native, zero-force structure, or, compare single-molecule force spectroscopy measurements with and without the putative allosteric regulator in the protein structure [18, 28, 29]. However, there are also situations where static mutated structures alone failed to explain the catch-bond behavior, as some of us recently demonstrated in the case of the FimH-manose interaction [30].

Rather than relying on indirect ways to correlate catch bonding behavior with different structural states, we want to directly predict the vinculin-actin catch-bonding behavior through atomistic molecular dynamics (MD) simulation techniques, which in principle allow us to predict the bound state lifetime using different forces simply by running many independent simulations and determining the average time to reach a dissociated state. Doing so would provide detailed molecular information about the possible unbinding pathways, and insight into which interactions are modulated by force, and how those contribute to the force dependence of the kinetics [5]. In practice, this is not easily achieved, because MD simulations of isolated proteins are currently computationally limited to the microsecond timescale, whereas the lifetime of protein-protein interactions are typically much longer (milliseconds to seconds) [5]; this problem is only exacerbated in the case of catch bonding behavior, since we are probing a situation where applying force could make the unbinding happen more slowly. Earlier studies of protein unfolding with force typically used non-physiologically large forces to observe significant changes within the accessible timescale of MD simulations [31–34]. Here, using these extra large forces would not test the fundamental hypothesis that small forces increase unbinding times.

To overcome these limitations, we deploy a sophisticated simulation strategy combining the application of directional constant pulling forces with *enhanced sampling* MD techniques; this allows us to compute the effect of applied forces on both estimates of unbinding free energy barriers and directly on unbinding rate constants for the interaction of the vinculin tail (Vt) with actin in a very large but still accessible amount of simulation time (∼ 200 *µ*s of MD for 250-300K atoms). While the combination of low forces and free-energy landscape determination using enhanced sampling has been done before for unfolding of coarse-grained proteins and small peptides [35–38], this has to the best of our knowledge never been deployed for such a complex and challenging system. We use this framework to elucidate and provide a molecular picture supporting models for directional catch-bonding underlying the Vt-actin interaction observed in single molecules studies.

## 2 Computational approach

### 2.1 Systems to study

It has been proposed that Vt’s catch bonding behavior arises when mechanical force promotes transition of the protein’s structure from a strongly-to weakly-bound state [39, 40]. In solution, the Vt protein consists of a bundle of five *α*−helices [41]. However, the CryoEM structure of Vt bound to actin showed a four helix bundle, with the fifth helix (H1) being unresolved, and hence this has been proposed to represent the strongly bound state [39]. It has been suggested that the weakly-bound state consists of a five-helix bundle similar to what is observed for Vt alone in solution; hence the ‘coupled folding-binding’ of H1 serves as an allosteric regulator of Vt affinity for actin [39, 40, 42]. For this mechanism, pulling on H1 in one direction versus the other has an asymmetric effect because one direction promotes dissociation while the other favors association [40]. *However, this does not fully explain why catch bonding behavior is observed in both directions, and which molecular motifs cause one state to be a strong binder and the other one a weak one*. Hence, here we wished to test whether Vt alone in either its weakly bound state or especially its strongly bound state could exhibit catch bonding behavior.

To do so, we created two primary systems to study (Fig. 1A,B). We first focused on the proposed strongly bound state which we termed ‘Holo,’ which is derived directly from the bound structure in Ref. 39 (PDB: 6UPW). Our second system, termed ‘Aligned’, was built by taking the isolated structure of Vt from Ref. 41 (PDB: 1QKR) and aligning the *α*-carbons of residues in helices H2-H5 with the structure in 6UPW. These two systems, which exhibits stable positioning and topology, serve as our model of the weakly- and strongly-bound states (Fig. S1–S4). Finally, we built a ‘Holo+H1’ system where the H1 residues were added to our equilibrated Holo structure in a disordered arrangement. Further details for all of these systems are given in the Methods Sec. 5.1 and simulation details are given in Sec. 5.2.

**Fig. 1.**
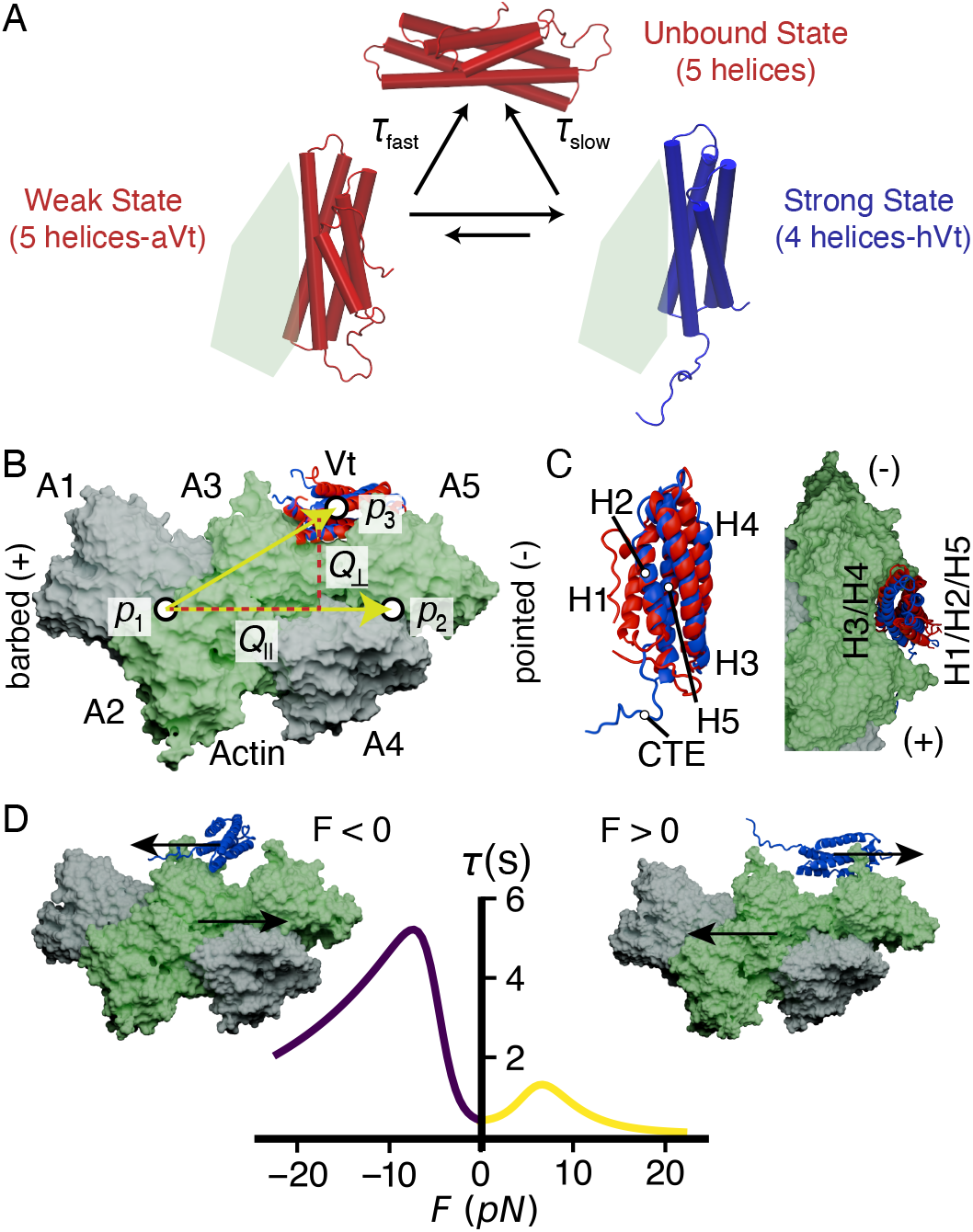
(A) Illustration of the two systems under study. It is hypothesized that upon binding actin (light green) the vinculin tail domain (Vt) converts from a five helical state to a four helical state, with the H1 helix being unresolved in the experimental structure. (B) Overlay of two Actin-Vt models. The Aligned (aVt, colored red) configuration contains helices H1-H5 while Holo (hVt, blue) has only H2-H5. Actin subunits A1 and A4 (colored gray) are restrained in the MD simulation to maintain the helical structure and prevent rotation. Arrows show primary collective variables (CVs) studied, Q_∥_ and Q_⊥_, which characterize movement of vinculin along and perpendicular to the filament, respectively. These CVs are defined using the vectors 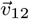 and 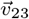 which point between the of center of mass (COM) of Vt helices H2-H5 (p_1_), the COM of actins A1 and A2 (p_2_), and the COM of actins A4 and A5 (p_3_). (C) Superposition of the Aligned and Holo configurations in an equilibrated configuration shown separately and as in their binding pocket, with α-helices and C-terminal extension (CTE) labeled. (D) Plot shows approximate experimentally derived Vt unbinding lifetimes as a function of force produced by a three-state model reported in Ref. 17 (see Sec. S1). Catch bonding behavior is observed when Vt is pulled parallel to actin in both directions, but is stronger when actin is pulled towards the pointed (−) end, i.e. Vt moves towards the barbed (+) end. Structures show representative snapshots from MD simulations combining pulling and enhanced sampling described below.

In all cases, the actin filament consists of 5 subunits. In order to roughly maintain the helical pitch of actin without directly effecting the interaction with vinculin, we add position restraints to the first and fourth actin monomers in the structure. This also allows us to use an oblong simulation box to minimize computational cost, since the filament cannot rotate. The filament is solvated with at least 1 nm layer of water, which we determined is sufficient to allow vinculin to fully unbind. We note that prior MD pulling studies on the Vt/actin interaction have used three actin [43, 44] based on the structure in Ref. 42, however using only three actin subunits requires that they be fully restrained, which we wanted to avoid as much as possible. Although we would like to do studies with even longer actin filaments to further minimize the effect of restraining part of the filament, the increased computational cost precluded doing so.

### 2.2 Pulling and biasing scheme

To pull Vt in a manner similar to single molecule experiments, we had to define two collective variables (CVs) that describe directions perpendicular and parallel to the filament respectively, *Q*_⊥_ and *Q*_∥_ (Fig. 1C, Sec. 5.1). Although large values of *Q*_⊥_ correspond to unbound poses of Vt, we also monitor the unbinding process through computing the fraction of key contacts maintained between H4 and H5 of Vt and actins A3 and A5 (*Q*_contact_) (see Sec. 5.3).

To probe the catch bonding behavior seen in the SM pulling experiments, a constant force is applied to *Q*_∥_ in either the positive or negative direction as oriented in Fig. 1. Experimentally, pulling actin towards its “pointed” end results in a stronger bond than towards its “barbed” end (Fig. 1D) [17]. To match this sign convention, we will define *F*_∥_ *<* 0 to be the direction of the stronger catch bond, where Vt moves towards the barbed end (Fig. 1D, inset).

We estimate the FES for Vt bound to actin using the On-the-fly Probability Enhanced Sampling/Meta-dynamics approach (OPES-MetaD) [45]. OPES-MetaD accelerates motion along chosen CVs by iteratively updating an applied bias that pushes the system away from previously explored regions (see Sec. 5.4). To probe the effect of force on Vt unbinding, we compute the FES in the two-dimensional space of *Q*_∥_ and *Q*_⊥_, while varying an additional constant force on *Q*_∥_ (or *Q*_⊥_). Because of the complexity of the unbinding reaction, we performed many independent OPES-MetaD simulations and then combined the separate estimates of the FES as described in Sec. 5.5. We emphasize that the OPES-MetaD bias is crucial in allowing the Vt protein to explore possible bound and unbound configurations on a sub-microsecond timescale, and otherwise it would be impossible to observe the effect of such small forces.

Similar biasing approaches can also be used to compute unbinding lifetimes [46]. We previously demonstrated that the force-dependence of unbinding rate constants could be captured through the infrequent Meta-dynamics (iMetaD) approach [46, 47], including catch bonding behavior for a model system, and subsequently we improved upon iMetaD estimates using a time-dependent rate formulation which performs better when non-ideal biasing coordinates are used [48]. Despite those improvements, iMetaD can break down for very complex unbinding problems, because it is difficult to prevent bias from being added during the crossing of the transition state. For this study, we adopted the related OPES-flooding approach (see Methods, Sec. 5.6), because it is possible to set a maximum amount of bias to be added and a region outside of which bias is not added, both of which help satisfy the assumptions underlying the kinetics calculations [49].

## 3 Results and Discussion

### 3.1 FESs differentiate proposed strong and weakly bound state

The FES projected along the two directions *Q*_∥_ and *Q*_⊥_, shown in Fig. 2, show clear differences between the Holo and Aligned models. We compute unbinding pathways by finding the minimum free energy path for unbinding using the string method [50], which are shown on top of the two dimensional surfaces. From the one-dimensional projections along the strings shown in Fig. 2C, we see that the Aligned state has approximately a 3.5 kcal/-mol smaller barrier to unbinding (Δ*A*^‡^) and a 2 kcal/mol smaller free energy difference with the unbound state, meaning we would predict that it is both kinetically and thermodynamically weaker than the Holo state. Results are quantitatively similar when projecting another set of coordinates Q_⊥_ and Q_contact_ (Fig. S5). The weaker starting state for Aligned leads to a different unbinding mechanism on average, where Aligned Vt tends to escape towards the barbed end of actin while the Holo Vt tends to escape towards the pointed end, which is the the direction of the smaller catch bond in experiment (Fig. 2A,B). Overall, these results confirm that model setups for Holo and Aligned can qualitatively capture the stability of the Holo state and the destabilizing effect of presence of the folded H1 helix in the Aligned state. These approximate one-dimensional free energy pathways also give us a way to define when the system has crossed into the unfolded state, and the values given in the inset of Fig. 2C will inform our later kinetics calculations. Further discussion of the molecular level differences between these two states is provided in Sec. 3.5.

**Fig. 2.**
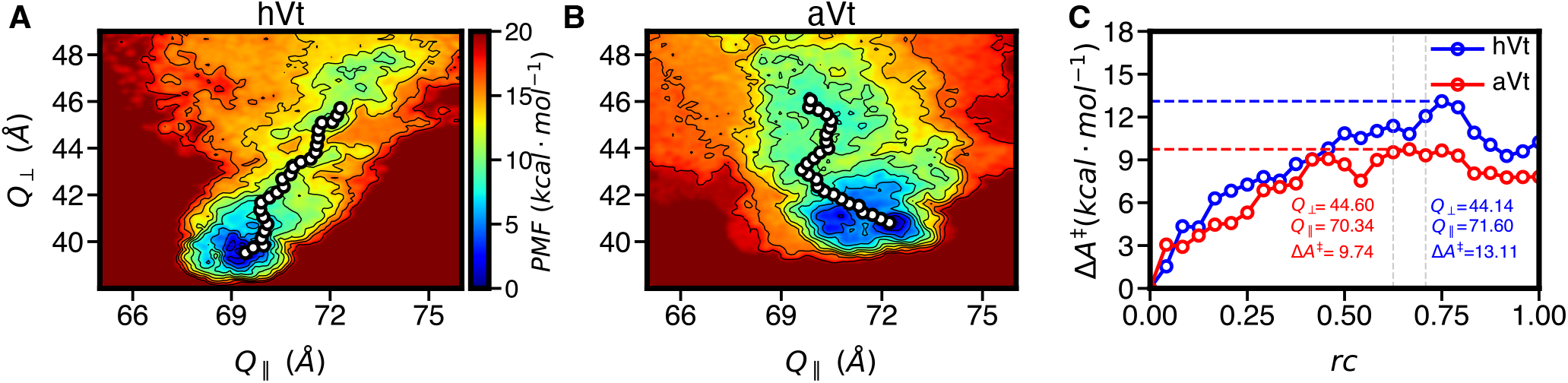
(A) FES for the Holo system (hVt) computed by 2D OPES-MetaD biasing, Q_∥_ and Q_⊥_. String shows the minimum free energy path on this surface going from the bound to unbound state. (B) Same as in A, for the Aligned (aVt) system. (C) One dimensional projections of the minimum free energy paths from A and B show that hVt is more stable and has a higher barrier to unbinding relative to Aligned. The Q_⊥_ and Q_∥_ values at the putative transition state are labeled; gray lines indicate points before the transition state and are used subsequently to help define the excluded region in unbinding rate calculations.

### 3.2 FESs with applied force predict asymmetric catch bond for strongly bound state and unidirectional catch bonding for weakly bound state

Having developed our protocol for computing the FES for Vt unbinding at zero force, we proceeded to repeat this procedure while applying physiologically relevant forces ranging from −30 pN to 30 pN along *Q*_∥_ as defined in Fig. 1B. We illustrate the effect of applied pulling force in Fig. 3A, where we see that the FES is tilted in the direction of the force for both Holo and Aligned systems. Typical unbound poses with force are shown in Fig. 3B. While in this figure we have only shown the effect of the largest and smallest forces, Fig. S6 shows results for all conditions.

**Fig. 3.**
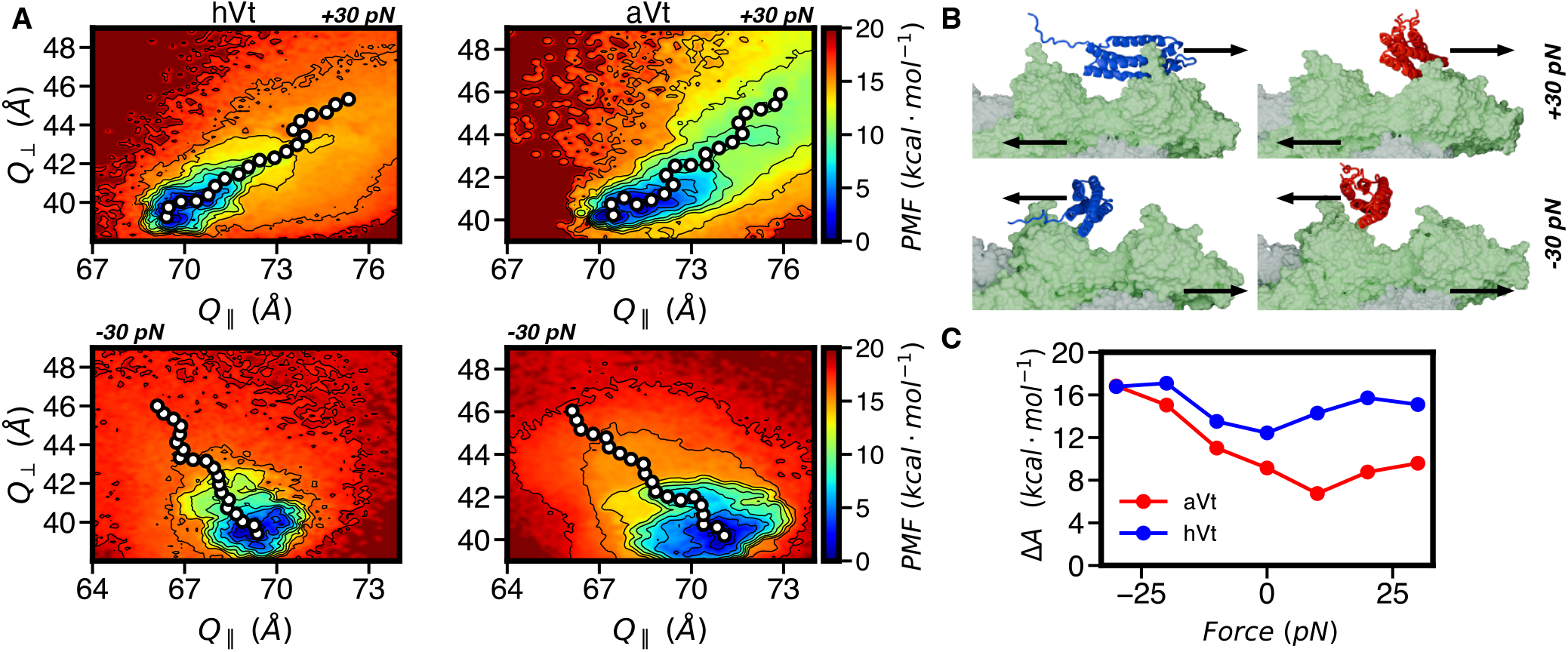
(A) 2D FES for Holo and Aligned as shown in Fig. 2, now with applied forces along Q_∥_ of +/- 30 pN, illustrating how applied force tilts the FES to larger or smaller values of Q_∥_. (B) Final snapshots of unbound states produced by positive and negative pulling forces of magnitude 30 pN along Q_∥_. (C) Transition barriers ΔA^‡^ along minimum free energy paths for Aligned and Holo. Holo shows an increase in barrier in both directions with force, with a slightly larger increase for negative F, while Aligned shows an increase only for negative F.

From the 2D FES in Fig. S6 we can compute 1D FESs by finding the minimum free energy paths as in Sec. 3.1 (Fig. S7) or by projecting onto *Q*_∥_ or *Q*_⊥_ (Fig. S8). In Fig. 3C we report the transition barriers (Δ*A*^‡^) extracted from the strings computed at different applied forces in both states. These data reflect a predicted catch bond for Holo, with the barrier increasing by 4 kcal/mol for negative forces and 3 kcal/mol for positive forces. In contrast, the Aligned state, which has a lower barrier in all but one case, shows catch bonding in the negative pulling direction but little change in the positive direction. The same conclusions can be reached if looking at the barriers in the curves of *A*(*Q*_⊥_, *F*) shown in Fig. S8.

This result is an indication that the different bound states of Vt may hold intrinsic catch bonding behavior, meaning not all of the catch bond need come from the transition between weak and strongly bounds states (see also [44]). We ascribe this force-induced increase in barrier to steric effects resulting from additional contacts with actin, which is why it can arise for either the weaker or strongly bound state. Further discussion of the molecular mechanism are discussed in Sec. 3.5.

Finally, we note that these results are suggestive, they do not quantitatively explain the observed experimental results, since the experimental changes in lifetime shown in Fig. 1D reflect a net 1.4 kcal/mol change in barrier in the negative direction, and 0.6 kcal/mol in the positive direction (if one assumes a constant prefactor the kinetic rate constant). This discrepancy could be due to the limitations of projecting a multidimensional FE landscape onto one or two dimension, as well as significant change in the position dependent friction (especially in light of the steric effects observed upon force application). Rather than only relying on our approximate approach to compute the FES, we also wished to directly estimate the lifetimes of the Vt/actin interaction at different applied forces.

### 3.3 Direct computation of unbinding times predicts intrinsic catch bonding in weak and strong states

We computed the lifetime of the Vt/actin interaction for both Holo and Aligned models for forces ranging from − 40 pN to 40 pN along *Q*_∥_ using the OPES-flooding approach (see Sec. 5.6 for details). We first calibrated our approach by performing 30 biased simulations at *F* = 0 for both models. We chose initial values of parameters defining when Vt is unbound and the maximum bias level parameters based on earlier iterations of the one-dimensional minimum free energy paths in Fig. 2C. Our bond lifetime calculations proved sensitive to the parameters Δ*E*, which is the maximum level of bias applied by OPES We observed that when the maximum bias level Δ*E* was above the height of the Holo barrier 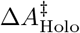, the predicted bond lifetimes were far longer than experiment. However, preliminary simulations with Δ*E* near the levels shown in Fig. 2C gave bond lifetimes on the same order of magnitude as experiments, which allowed us to settle on a value of Δ*E* = 12 kcal/mol for Holo and Δ*E* = 10 kcal/mol for Aligned and an excluded region *Q*_⊥_ > 44 Å.

With these OPES-Flooding parameters, we computed a bond lifetime of 2.3 *±* 0.37 sec for the Holo state and 0.13 *±* 0.02 sec for the Aligned state (Fig. 4). Although Holo and especially Aligned are just models for the weak and strongly-bound states, these lifetimes are physically reasonable, and in particular, the lifetimes for Holo are well within the range of lifetimes observed from individual measurements [17]. This correspondence was closer than we would have predicted for OPES-flooding, and we therefore considered our choice of CVs and parameters of sufficient quality to consider assessing the lifetime at different forces.

**Fig. 4.**
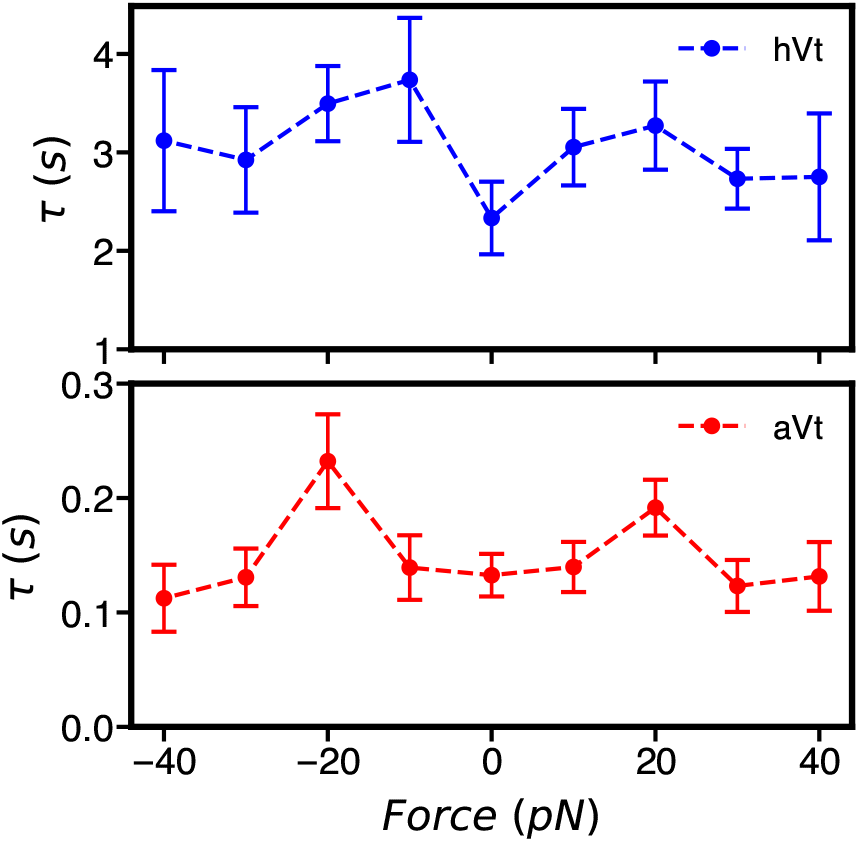
Lifetimes of Vt unbinding extracted by OPES-Flooding for Holo (top) and (Aligned) bottom. Both show minimal sensitivity to applied force along Q_∥_ in this physiological force regime. These are obtained by fitting cumulative distribution functions shown in Fig. S9. Error bars are from bootstrapping as described in Sec. 5.6.

To compute the rates with different applied forces along *Q*_∥_, we repeated these same calculations, performing simulations launched from at least twenty starting points in the equilibration trajectory at each force (see Sec. 5.6). In ideal circumstances, one might want to choose different OPES-Flooding parameters for each force; but, for these simulations we elected to use the same biasing parameters for all forces. This was done both for simplicity and to avoid implicitly selecting parameters that favored a hypothesis of how the system should behave.

As seen in Fig. 4, the lifetimes at different forces for Holo and Aligned both exhibit catch bonding behavior within this force range, with longer lifetimes under applied force than at zero force. This is in general accord with our earlier FES calculations (Fig. 3C), but with less than a two-fold increase for both states. Combining our FES and unbinding lifetime calculations, we would conclude that there is some intrinsic catch bonding property that arises when pulling laterally on actin due to the increased interactions with actin under force (see Sec. 3.5). Our computed lifetimes depend assymetrically on force, and they do suggest a stronger catch bonding effect in the negative force direction as observed in experiment, although to a lesser degree [17].. At the largest forces assessed here, the lifetimes approximate decrease back near the zero force value, rather than exhibiting a an exponential slip bonding decrease. This behavior is compatible with the mild force-dependence observed experimentally for the supposed Aligned state (i.e., at positive force in [17]) as well for the an artificial Holo construct of catenin [18] missing the H1 helix (and thus analogous to our own Vt construct here).

While the magnitude of the intrinsic catch bond effect we predict here is smaller than the total observed effect in experiment, intrinsic catch bonding in the strong and/or weak state is also compatible with the model proposed wherein a dominant source of catch bonding is the transition between Holo and Aligned states, as discussed more in the next section. Given our data in Fig. 4, the *maximum* catch bond that could be obtained would be if pulling actin towards the pointed end completely promoted a transition from a stable Aligned state (at 0 pN, *τ* ∼0.13 sec) to the Holo (at −10 pN, *τ* ∼ 3.8 sec) state, which would result in a 30-fold effect; this is larger than the 10-fold effect measured for Vt (Fig. 1D) [17].

### 3.4 N-terminal pulling can trigger motion from Holo towards Aligned

Experimental unbinding lifetimes in Ref. 17 were determined by an optical trap (OT) based assay where the vinculin head (Vh) domain is bound to a platform bead allowing Vt to bind to actin held taught by two microspheres. When actin is pulled in either direction, tension is introduced onto the actin/Vt interface via pulling on Vt’s N-terminus, close to the region containing helix H1 in our Aligned structure. As discussed above, it has been proposed that pulling forces asymmetrically promotes unbinding of H1 due to pulling on its N-terminus, though we sought to test this hypothesis using our MD setup.

To do so, we employed our Holo+H1 model, which had the H1 helix grafted to our Holo structure random orientation, dissociate from H2-H5, as shown in Fig. 5A. To apply pulling forces, we redefined our CVs *Q*_∥_ and *Q*_⊥_ using the COM of the final ten alpha carbons of the N-terminus of Vt rather than the COM of H2-H5. We performed five independent equilibrium MD simulations at 50pN and −50pN along *Q*_∥_ to observe the behavior of the *N*-terminus.

**Fig. 5.**
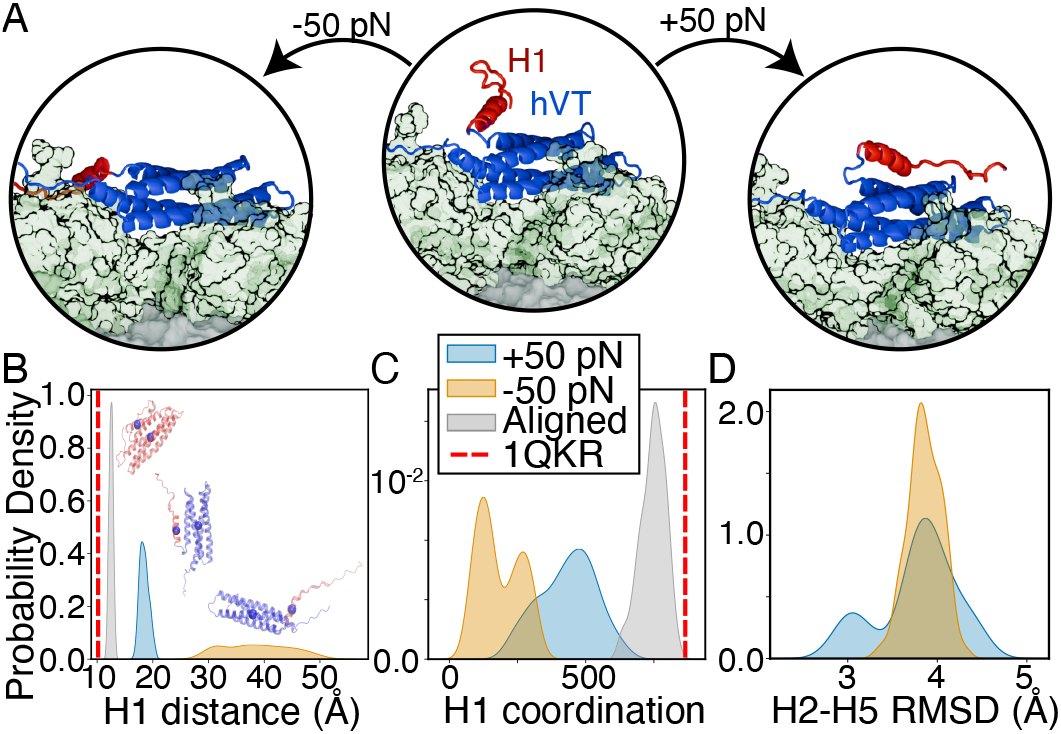
(A) Snapshots showing the initial configuration of our Holo+H1 system, and final configurations after pulling on the N-terminus at *±*50 pN along Q_∥_. Pulling the N-term towards the pointed end (+ 50 pN) docks H1 along the bundle, while pulling towards the barbed end (−50 pN) extends H1 away from the rest of Vt. (B) Histogram of the distance between the COM of H1 and the COM of H2-H5 under force. The Aligned system (gray) is shown in gray and maintains a distance close to the crystal structure (dashed line), while pulling with positive force (blue) and negative force (orange) result in more or less association, respectively. (C) Histogram of the number of contacts (atom pairs with distance below 5 Å) between H1 and H2-H5 using atoms carbon oxygen and nitrogen. (D) Histogram of the C_*α*_ RMSD between helices H2-H5 in Holo+H1 and H2-H5 in 1QKR (1QKR).

We find that when the N-terminal of Vt is pulled toward the pointed end of actin, H1 can be extended away from the H2-H5 bundle. As a consequence, it ends up forming little to no interaction with the rest of Vt (Fig. 5B and C). In contrast, when pulled toward the barbed end, it closely interacts with the rest of the bundle (Fig. 5B and C), forming a large fraction of the number of interactions present in the aligned state. Noticeably, in two out of five simulations the configuration of H2-H5 spontaneously becomes more similar to the crystal structure of Vt unbound, and hence shifts towards our model of the weak binding state, and this is reflected in the histogram of RMSDs shown in Fig. 5D. Further enhanced sampling studies probing the transition between the structure of the Holo and Aligned H2-H5 arrangements would be needed to confirm this result (see Conclusions).

In summary, pulling the N-terminus of Vt towards the barbed end and actin towards the pointed end (*F <* 0) disfavors the five-helical weak binding state, whereas pulling with *F >* 0 either promotes or is at least commensurate with the proposed weak binding state. Hence, these simulations thus provide evidence supporting the model proposed in Ref. 40, where forces asymmetrically contribute to the removal of H1 from the rest of Vt. However, this does not fully explain why Vt can exhibit catch bonding behavior when actin is pulled in either direction. In the next section, we discuss the major differences between our Holo and Aligned model and changes induced by force.

### 3.5 Molecular insight into Vt catch bond

Above, we demonstrated that our Holo model derived from the bound Vt structure in Ref. 39 and Aligned model derived by combining the structure in Ref. 39 with the crystal structure from Ref. 41 are both stable and have unbinding lifetimes close to what would be expected from single molecule experiments. In Fig. 6A and B, we show representative snapshots (see Sec. 5.7 for how these were chosen) from 500 ns of equilibrium MD for both systems; using these data we are able to highlight some important differences between the states that contribute to the difference in binding stability of our Holo and Aligned models.

**Fig. 6.**
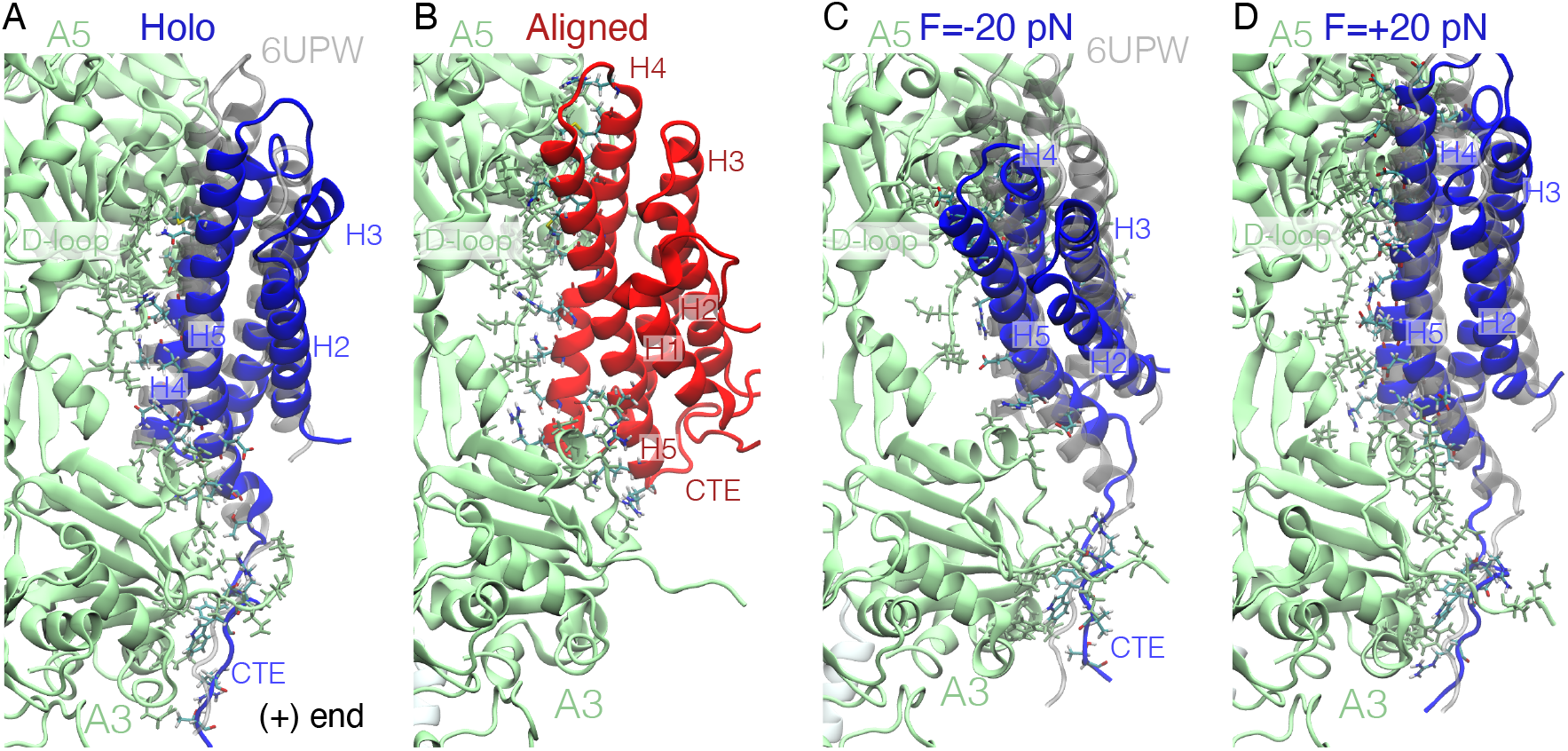
Representative structures of Vt bound to actin, with hVt in blue, aVt in red, and actin in green. The initial structure (6UPW) is overlain on the Holo structures after superimposing actin A3 from the MD and EM structures. Residues in Vt and actin A3 and A5 separated by less than 3 Åare shown in VMD’s [51] ‘licorice’ representation. (A,B) Representative structures from 500 ns of equilibrium MD on Holo and Aligned systems. (C,D) Representative structures from OPES-MetaD simulations of Holo with and without force illustrate the primary changes induced by pulling on Vt’s COM.

From our calculations, the Holo or putative strong binding state is on average bound closer to the center of the actin filament, makes more contacts with actin, has more buried surface area (BSA), and has a stronger energetic interaction association (Fig. S1–S4 and Fig. S10). However, when decomposing these effects into contributions from different parts of Vt, we find that a major contribution to the number of contacts, buried surface area, and energetic attraction in Holo comes from a disordered C-terminal extension (CTE), and those quantities are relatively similar when comparing only H2-H5 between Aligned and Holo (see Figs. S2 and S3). This is in accordance with earlier hypotheses that the CTE may play a major role in stabilizing interactions of Vt with actin [39, 40], as well as in detecting stretched states of actin [39]. We note that the CTE in the Aligned model stays bound on the opposite side of the bundle within the timescale of our simulations, and as discussed in the Conclusions, separate investigations would be needed to probe whether an intermediate Aligned-like state with extended CTE is stable.

Given that there are a similar number of contacts and buried surface area in H2-H5 between Aligned and Holo, we also considered how these contacts are distributed. In Fig. S11 we highlight the residues in close contact in each state in the representative structures. Our observations suggest that the Aligned or putative weak-binding state involves a tilt of the bundle such that aVt’s longer H5 helix is able to take up most of the contacts with actin; in contrast, contacts between hVt and actin in the Holo state are more evenly distributed across H4 and H5 helices, and this may confer some additional kinetic stability. We also observe in the Aligned state an extension of actin A3’s D-loop, which allows it to maintain contacts with Vt.

We are also able to use the molecular data from our extensive OPES-MetaD simulations to investigate the effect of force on the bound pose of Vt. OPES-MetaD produces biased sampling of structures, meaning that we cannot simply look at the output configurations to infer mechanistic details. However, we have already ascribed a biasing weight per snapshot produced from these simulations (see Sec. 5.4, 5.5); these weights are exponentially larger for low free-energy states as compared to those in the barrier or unbound states. Hence we can compute weighted averages of molecular quantities and use our approach from Ref. 52 to select representative equilibrium frames which reflect information about the bound structural ensembles.

In Fig. 6C and D we show representative structures in the Holo state with force applied to *Q*_∥_, towards and away from actin’s barbed/pointed ends (see Sec. 5.7 for details). When negative force is applied, meaning that actin is pulled up and Vt down, on average it causes Vt to rotate in such a way that more contacts are formed between actin A5 and the bottom/middle of H4, and contacts on H5 are decreased with A5 but maintained or increased with actin A3. At the same time, the CTE is extended, making additional contacts with actin A3. In contrast, when force is applied in the positive direction, it causes Vt on average to rotate in the other direction. Here, more contacts are formed between H4 and H5 with actin A5, but contacts with A3 appear somewhat decreased. To make these observations more quantitative, we compute the weighted histogram and weighted average amount of BSA as a function of applied force. As shown in Fig. S10, total BSA increases when pulling in the negative direction for Holo (and also Aligned), and it is basically maintained when pulling in the positive direction. We believe this is a major contributor to the predicted increase in unbinding barrier and bond lifetime with applied force(Fig. 3). The fact that contacts and buried surface area can be maintained when pulling in either direction due to the crescent shape of the actin side binding pocket helps explain why the unbinding rate does not increase even with relatively large applied forces.

## 4 Conclusion

In this work, we have probed the molecular mechanism of the catch bond formed between the vinculin tail domain and the cytoskeletal polymer actin using molecular dynamics simulations. Thanks to a combination of enhanced sampling techniques, we are able to address experimentally-relevant seconds timescales and physiological forces of a few pN on a large macromolecular assembly. This unique approach enables us to directly estimate the unbinding lifetimes of a catch bonding system at the molecular level, and to probe the force dependence of the barrier to unbinding for such a complex system. Moreover, our approach can be extended to probe the response to force for other analogous systems. Our results are in nearly quantitative agreement with experimental measurements, for which they provide a molecular interpretation. The atomistic insight gained from these studies can further guide future mutational studies to probe this catch bonding mechanism involved in mechanotransduction.

More specifically, we interrogated a three-state catch bond model, consisting of an unbound state, a weakly bound state and a strongly bound state. Through our MD computations, we established that both our models of the strongly-bound (built from the CryoEM structure of Vt bound to actin with an unresolved H1 region) and weakly-bound (built from a complete, isolated Vt structure) states, have binding lifetimes compatible with that predicted from single molecule studies. We predict that for forces around 10-20 pN it is possible for either the strongly- or weakly-bound structures to shift into configurations that take longer to unbind than at zero force (see also, e.g., Ref. [44]). However, this effect is not as large as what is observed in experiment, suggesting that the mechanism of catch bonding derives from a superposition of intrinsic catch bonding with the previously proposed three-state model of Vt’s catch bond, which involves allosteric regulation of binding affinity by the H1 helix. This puts a focus on whether pulling on the H1 helix gives a directional dependence to the catch bonding behavior, as pulling in the direction of the strong catch bond moves H1 away from Vt. Our MD simulations support this hypothesis, but further enhanced sampling simulations probing the free energetic barrier to transition between the weak and strongly bound configuration of H2-H5 while bound to actin at different forces would be important to further confirm this mechanism.

There are still inherent limitations to our approach. At relatively high forces (> 30 pN), our calculations predict lifetimes similar to zero force, whereas experimental measurements are indicative of a slip bond system in that regime. This might be a consequence of the direction of pulling and how we performed the pulling, namely pulling the COM of Vt directly parallel to the actin filament; this pulling direction moves Vt into direct steric contact with parts of actin’s side binding groove, and we often observe that Vt has to slide perpendicular to the filament (in and out of our diagrams) on its way to unbinding. The exact details of how the SM experiments were performed, in which Vt is attached to a surface via linkers attached to its N-terminus, and how actin was suspended between beads, may induce angular forces on the molecule and some tension the two non-actin-parallel directions; we would like to incorporate these aspects of the experiment into further investigations.

The apparent increase in barrier with applied force from our FES calculations is larger than what would be expected from experiments, and larger than what we would predict with our rate calculations. Our current feeling is that the kinetics calculations may more accurately reflect reality, since our FES calculations (a) were approximately computed from many short OPES-MetaD simulations and (b) inferring kinetic information from these involves looking at a 1-dimensional reaction coordinate, which could have a misleading barrier height as compared to the true barrier for the reaction would the exact reaction coordinate be known. Moreover, just looking at the barrier along our 1-d free energy surfaces does not take into account that there could be multiple reaction pathways, and that the diffusion constant along the reaction path might change with force. We would like to investigate this discrepancy in the future by considering previously proposed mutations that modify the Vt unbinding lifetime and also by considering an analogous protein like *α*-catenin to provide additional insight into the performance of our methods.

In the future, we would also like to try alternative approaches to computing the minimum free energy pathway for unbinding; for example, the data we already generated could be a good starting point for performing variations of the string method in high dimensional space [53]. We can also use the large amount of unbinding trajectories we have generated to produce machine learned reaction coordinates that better describe the unbinding mechanism, which could lead to both better FES and rate calculations, as well as additional insight into the true unbinding mechanism [54, 55]. Finally, we have not considered an intermediate between our Aligned and Holo model, which has a five-helix bundle state but CTE bound to actin, and it would be intriguing to investigate whether this is a stable structure and if it plays any role in the overall mechanism.

## 5 Methods

Below we describe preparation of our systems, MD simulation details, and enhanced sampling methods. System inputs and simulation parameter files, analysis code, and equilibrated system structure files are available from https://github.com/hocky-research-group/Pena-Vinculin-Unbinding.

### 5.1 System preparation and nomenclature

The structure 6UPW [39] consists of five actin subunits A1-A5 and two bound (meta)Vinculin tail domains [39]. An initial template system was prepared by removing the Vt bound between actin subunits A1 and A3 in 6UPW, leaving one Vt bound between A3 and A5. Missing residues (1-4, 375) in actin subunits were included using VMD’s modelmaker function [51]. Actin subunits contain an ADP, Mg^2+^ ion, and water [56]. This structure was used as a starting point to build all models.

The Holo system was constructed by solvating the reference structure with TIP3 water, and then 50 mM of NaCl was added in addition to neutralizing with sodium ions. The hVt protein in this system consisting of four helices (H2-H5 bundle) and a CTE tail. hVt has residue ids: 981-1131 (151 residues).

The Aligned system was constructed by first aligning the H2-H5 C_*α*_ atoms in Ref. [41] to the H2-H5 bundle of Vt in the reference structure with UCSF Chimera’s matchmaker program [57]. The resulting structure was later solvated and ionized as in Holo. aVt has residue ids: 882-1061 (180 residues)

The Holo+H1 system was built by positioning H1 of aVt near H2 of hVt and a bond was created manually between the last H1 and first H2 residue with Chimera, followed by solvation and addition of ions as before. The nhVt proteins consists of five helices, H1-H5 with H2-H5 in a bundle and a CTE tail. nhVt has residue ids: H1 882-912, and H2-H5+CTE 981-1131 (182 total residues). We initially used a minimum padding of 1.2 nm of water in all directions, and this Aligned and Holo box size was used for the FES calculations. After determining that a smaller box was sufficient to observe unbinding, we created smaller versions of the same system with a 1 nm minimum padding, which allowed us to perform our rates calculations more efficiently.

### 5.2 Simulation details

Minimization and equilibration simulations were performed in GROMACS-2023 [58], and all production simulations as well as post-processing were performed with GROMACS-2023 patched with PLUMED versions 2.8 and 2.9 [59]. The CHARMM36 [60] force field was used for all the bonded and nonbonded parameters of proteins. Equilibration was performed in a manner similar to Ref. 61. For FES calculations under mechanical load, 20 independent minimization and equilibrations were performed. Configurations were minimized with the steepest descent algorithm for 5000 steps with *dt* =1 fs followed by a 5 ns constant-volume and temperature (NVT) equilibration stage (2 fs timestep) where A1, A4 and Vt components had position restraints of force constants 1000 kJ/(mol nm^2^) on backbone heavy atoms to allow the solvent other components to relax. Furthermore, protein and solvent atoms were coupled separately to a 300K temperature bath using GROMACS’s v-rescale thermostat. This was followed by a 5 ns equilibration (2 fs timestep) at constant temperature, a constant pressure (NPT) using GROMACS’s C-rescale barostat with 1.0 bar reference pressure with position restraints on Vt still on. An additional 5 ns NPT equilibration using the nose-hoover thermostat was performed with restraints on and a final 5 ns NPT equilibration was carried out this time using GROMACS’s Parinello-Rahman barostat without restraints on Vt. For productions runs, positions restraints were kept on for A1 and A4, which maintained restraints on the backbone heavy atoms using a force constant of 1000 kJ/(mol nm^2^).

For the 500 ns long equilibration runs, we followed the same steps as above for initial equilibration but kept the V-rescale thermostat and the Parinello-Rahman barostat combination for the long 500 ns run. For rate calculations, starting configurations were extracted from the latter half of the 500 ns trajectory for each model and each configuration was equilibrated for 5 ns under NPT using the the v-rescale thermostat and c-rescale barostat with restraints on Vt and 5 additional ns under NPT using the parrinello-rahman barostat and without restraints on Vt. Resulting structures were used in production runs under NPT.

For all equilibration and production runs, long-range electrostatics were calculated using the Particle Mesh Ewald (PME) algorithm with the cut-off for short-range non-bonded interactions at 1.2 nm. Bonds between hydrogen and heavy atoms were constrained using the LINCS algorithm. Finally, the equations of motion were integrated every 2 fs using the Velocity-Verlet algorithm (see Sec. 5, Fig. S1).

### 5.3 Key collective variables

To characterize the motion of Vt relative to actin, we used PLUMED [59] to define two vectors shown in Fig. 1B: one pointing from the center of A1 and A2 to the center of A3 and A4 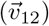 and one pointing from the center of actins A1 and A2 to Vt 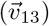. After defining these vectors, *Q*_∥_ is defined as the scalar projection of 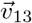 along 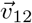 and *Q*_⊥_ is defined as the length of the perpendicular component of 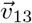 relative to 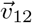. Although large values of *Q*_⊥_ correspond to unbound poses of Vt, we also monitor the unbinding process through computing the fraction of key contacts maintained between H4 and H5 of Vt and A3 and A5 (*Q*_contact_). In this work we define *Q*_contact_ as in Ref. 62, as a sum of fractional contacts *Q*_*ij*_ between key selected pairs of atoms where the *Q*_*ij*_ for each distance is computed as:

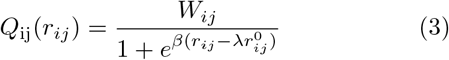

where *λ* is 1.8, *β* is 5.0 Å^−1^, *r*_0_ is 4.0 Å, *r* is the distance *i* at time t, and *W*_*ij*_ is the weight for *Q*_*ij*_ which is defined to be the constant 1*/N*_pairs_. Atom pairs are specified in table 2 in the SI.

### 5.4 The OPES-MetaD method

The on-the-fly probability enhanced sampling metadynamics (OPES-MetaD) method [45] was used to rapidly explore configurations of Vt relative to actin using the aforementioned CVs. This method is similar to metadynamics [63] however OPES-MetaD computes the bias from the ratio of a target probability distribution to the reweighted probability distribution of the chosen CVs, making the bias nearly constant as the system FES is refined (as compared to standard MetaD which continually adds Gaussian functions to the applied bias). The reweighted probability distribution is obtained via kernel density estimation and the bias on CVs **s** has the form:

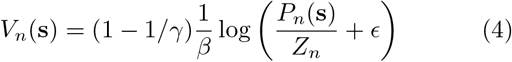

One particular parameter that provides an advantage over standard MetaD is the Δ*E* barrier parameter which is used to set the bias factor *γ* (which determines how much FES barriers are reduced) and *ϵ* in Eq. 4 as in *γ* = *β*Δ*E* and 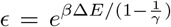. This barrier parameter in practice is chosen to be roughly equal to the energy barrier to be overcome during simulation. In essence we can limit the sampling of higher energy regions that may not be realistically accessible. OPES-MetaD also allows us to define an excluded region outside of which the bias goes to zero, which we use in our computation of binding lifetimes.

### 5.5 Computing the FES for Vt unbinding with OPES

In general, it is difficult to compute an unbinding FES even when using a biasing scheme such as OPES-MetaD because of the need to fully explore and observe many transitions between the bound and unbound state [64, 65]. Therefore, to compute an approximate FES which captures the bound state and transition to the barrier, we first ran 20 separate OPES-MetaD simulations for 100 ns with harmonic upper walls at *Q*_⊥_ > 52, *Q*_∥_ > 80 Å and a harmonic lower wall on *Q*_∥_ *<* 57 Å(positions taken from initial equilibration runs). The spring constant for the wall potentials was set to 1000 kcal/mol/Å in all cases. For FES calculations in the absence of force, we set Δ*E* to 25 kcal/mol, and then for those under load, we set Δ*E* to 20 kcal/mol. We then combined these independent sets of data using WHAM (as implemented in PLUMED [59]) to combine the final quasi-static bias from each different simulation to produce an FES estimate along the two biased CVs for both Aligned and Holo models (Fig. 2A,B at zero force and Fig. 3, S6 with applied forces). To do so, we used PLUMED to recompute what the bias would have been in trajectory *i* if we had used the OPES bias from simulation *j*. We then computed the minimum free energy path on each surface by running the string method on the potential [50]. This allows us to define a predicted barrier height and to define *Q*_⊥_ values beyond which we consider Vt to be unbound (Fig. 2C). We also projected these CVs into the space of *Q*_⊥_ and *Q*_contact_ which allowed us to confirm the unbinding of the Vt (Fig. S5).

### 5.6 Rates of Vt unbinding with OPES-flooding

We computed the average times for unbinding using the OPES-flooding approach [49]. Based on the earlier InfrMetaD [46] and hyperdynamics [66] approaches, OPES-flooding assumes that the process to be studied is a rare event characterized by a large free energy barrier. It then seeks to apply a biasing potential only in the starting state without applying bias to the system as it crosses the transition region. In this case, the speedup or acceleration factor for simulation *i* can be computed as 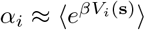, where the average is taken over the trajectory up to the point where the event occurs, *t*_*i*_. To compute the average unbinding time *τ*, we fit the cumulative distribution function of rescaled unbinding times (*T*_*i*_ = *α*_*i*_*t*_*i*_) to an exponential distribution [49, 67],

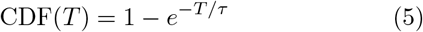

The goodness of fit can be checked by a Kolmogorov-Smirnoff (KS) test [67]. Error bars are computed via bootstrapping, where 200 sets of the same number of biased trajectories are selected with replacement, the CDF fit is performed, and then the standard deviation of fit *τ* s are computed [68].

As stated in the Results section, we used Δ*E* = 12 kcal/mol for Holo and Δ*E* = 10 for Aligned, and set an excluded region *Q*_⊥_ > 44 Å. We define an unbinding event when *Q*_⊥_ > 46 and *Q*_contact_ is below 0.25. We performed 30 OPES-flooding simulations at zero force and 20 for all other forces for each system until Vt was determined to unbind. ECDFs pass the KS goodnes of fit test in almost all cases suggesting we have a reasonable choice of CVs and excluded region for this setup S9.

### 5.7 Choosing representative configurations

To select representative frames for Fig. 6, we used the ShapeGMMtorch package from Ref. [69, 70]. To do so for equilibrium simulations in Fig. 6, we performed iterative alignment of the Vt structure using all backbone atoms to a mean structure taking into account the covariance of positions using the “Kronecker form” of the covariance matrix described in Ref. [69]. This approach naturally favors the regions which are less floppy. The result of this procedure is a single multivariate Gaussian in Cartesian position space, and we then selected the frame from the MD trajectory which had the highest likelihood in this distribution (having the minimum Mahalanobis distance from the mean).

We recently extended our ShapeGMM approach to MetaD/OPES simulations [52], meaning that we can take into account the weights of each frame in biased simulations and fit an equilibrium ShapeGMM model. To pick the structures in Fig. 6B, the same iterative alignment procedure was performed using C_*α*_ atoms of Vt, with weights of each frame given by 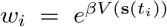, where **s** represents the position in *Q*_∥_ and *Q*_⊥_. This was per-formed for each OPES-MetaD trajectory at *F* = ± 20 pN separately (so that relative weights did not have to be determined), to obtain representative frames for each run, and then one was selected manually for each force.

## Acknowledgements

We thank Professors Gregory Alushin, Alexander Dunn, and Brenton Hoffman for helpful discussion about this system and experimental details. WJPC, FM, and GMH were supported by the National Institutes of Health through the award R35GM138312 (and R35GM138312-02S1 to WJPC). This work was supported in part through the NYU IT High Performance Computing resources, services, and staff expertise, and simulations were partially executed on resources supported by the Simons Center for Computational Physical Chemistry at NYU (Simons Foundation Grant No 839534). This work was supported by the “Initiative d’Excellence” program from the French State (Grant “DYNAMO”, ANR-11-LABX-0011-01 to GS).

## Supporting Information

### S1 The three state catch-bond kinetic model

The following describes the three state model from Ref. 17 (parameters in Tab. 1) which was used to fit the experimental data for Vt unbinding, and was used to produce the curve in Fig. 1D.

**Table 1.** Parameters for a catch bond model for Vt unbinding, taken from Ref. 17.

| rate | $k_{ij}^0$ ($s^{-1}$) | $x_{ij}$ | $x_{ij}$ |
| --- | --- | --- | --- |
| $k_{10}$ | 3.9 | 0 | 0.5 |
| $k_{20}$ | 0.045 | 0.3 | 0.7 |
| $k_{12}$ | 2.7 | 0 | 0 |
| $k_{21}$ | 8.3 | -3.8 | -2.8 |

**Table 2.**
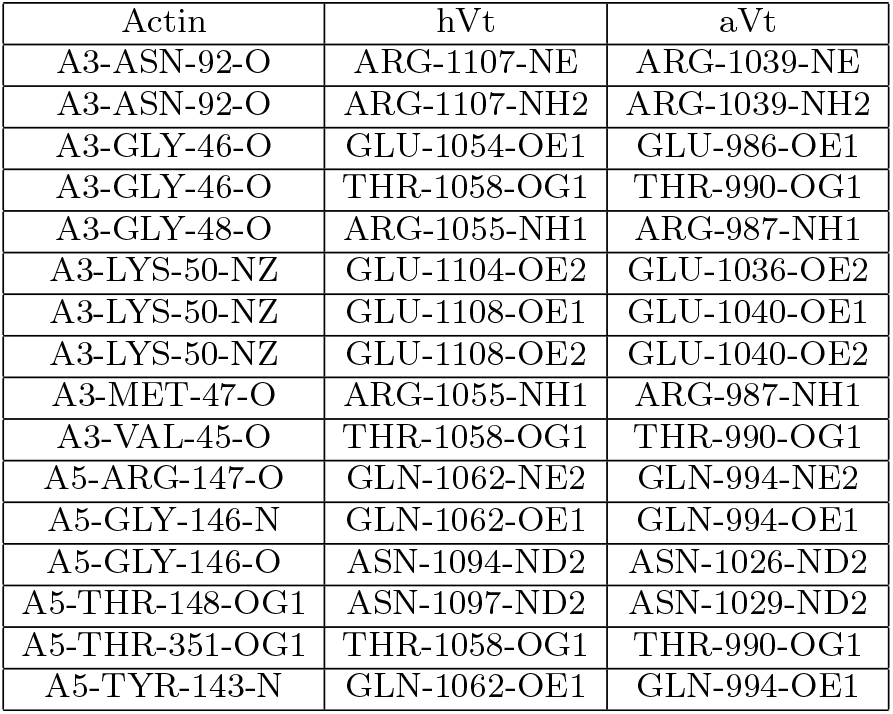
Atom pairs used in *Q*_contact_. Residue numbering is based on original residue numbers in the PDB structures.

In state 1, Vt is weakly bound and in state 2 Vt is strongly bound. Vt transitions between states 1 and state 2 with rates *k*_12_ and *k*_21_, respectively. In state 0, Vt is in the unbound state which can be reached from state 1 with rate *k*_10_ or from state 2 with rate *k*_20_.

The cumulative distribution function (CDF) for the three state catch-bond model is given by:

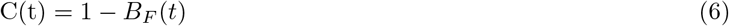

and its derivative (PDF) is given by:

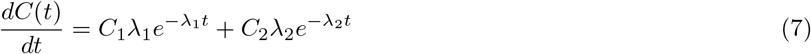

The parameters *C*_*i*_ and *λ*_*i*_ above were derived in [22, 71] as follows:

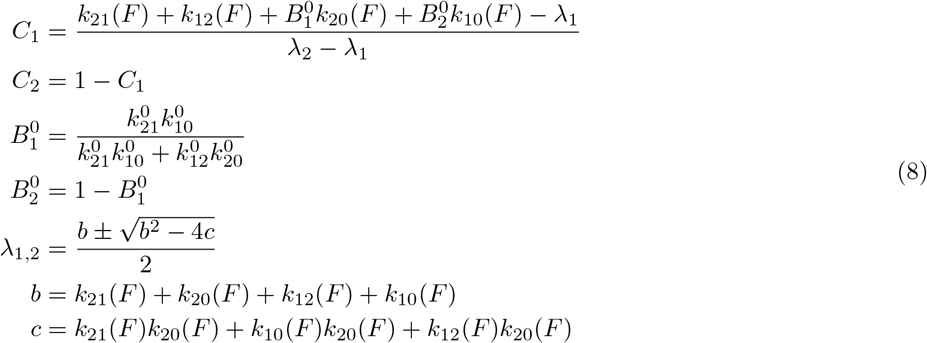

It was assumed that each individual transition from one state to another is described by Bell’s model [13]:

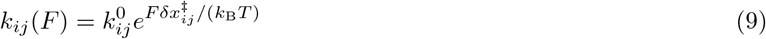

where 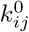 is the transition rate from state *i* to state *j* in the absence of force and *x*_*ij*_ is the distance of the transition state from state *i*.

Although generally unbinding events under load are well described by Bell’s model where the unbinding rate increases with increasing force (the exponent is positive), for the unbinding of Vt from F-Actin, the proposed mechanism in Ref. 17 is that the rate of transition from the strongly bound state to the weakly bound state decreases significantly under load (exponent is negative) and thus the overall unbinding rate decreases with increasing load. In this context where there is a transition from one bound state to the other, a more complex approach is required to obtain the mean transition lifetime from bound to unbound state as a function of force. In practice, the pulling forces and their corresponding set of unbinding times are used as inputs to optimize all 8 parameters: *k*_*ij*_ and *x*_*ij*_ via maximum likelihood estimation (MLE). The objective function is the bond survival function described by a double exponential decay. The probability distribution function of bond lifetimes can then be computed as the derivative of the cumulative distribution function. From this result, the mean lifetime of the transition at each force can be calculated as in Eq. 10 [22, 71]:

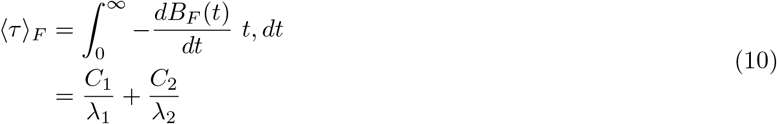

where ⟨*τ*⟩ _*F*_ is the mean lifetime of the bond (*k*_*F*_ = 1*/* ⟨*τ*⟩ _*F*_) and *B*_*F*_ (*t*) is the bond survival function given by a double exponential decay:

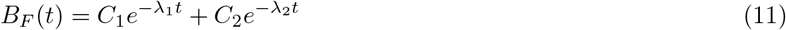

### S2 Additional figures

**Fig. S1.**
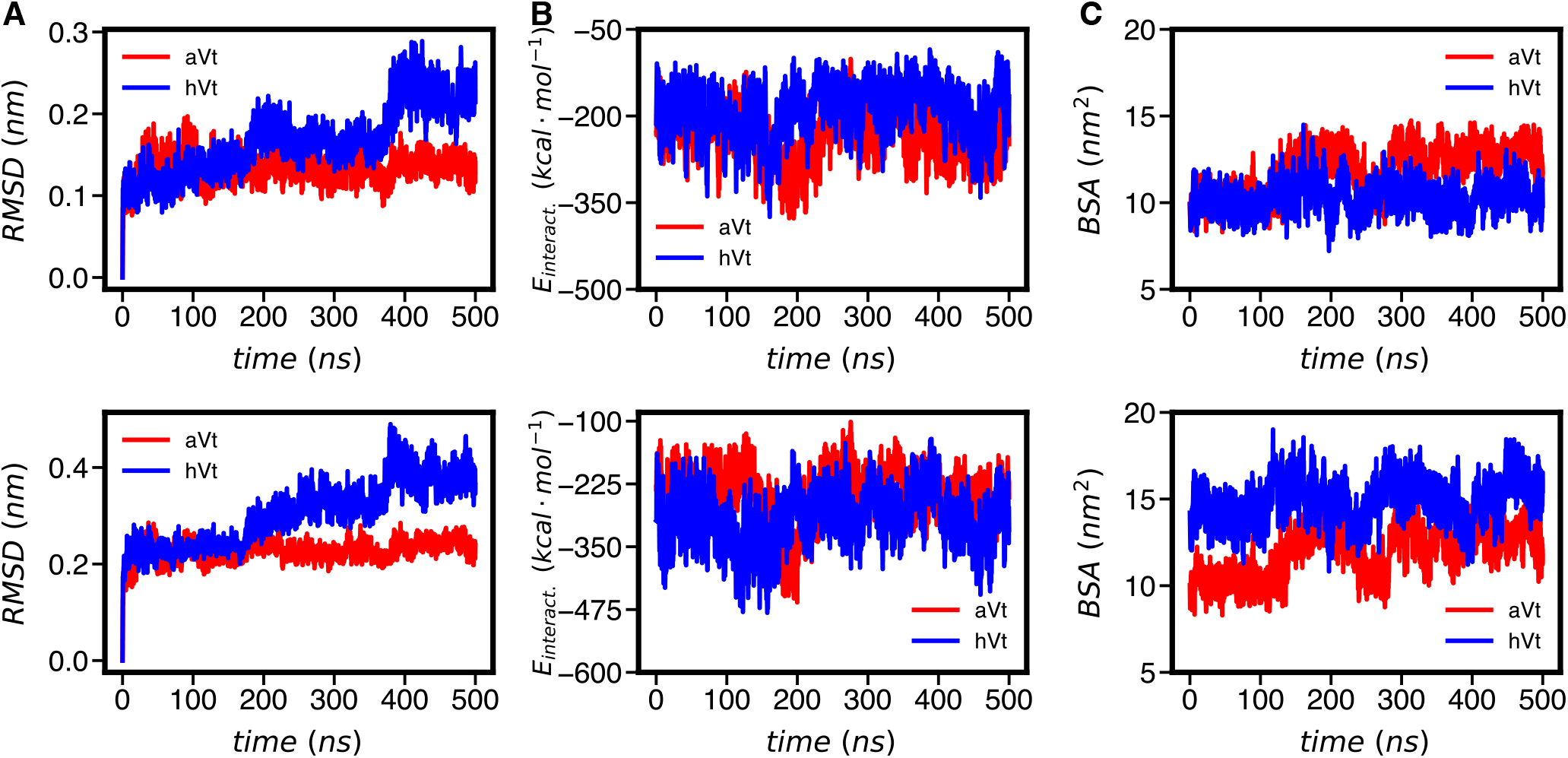
Top figures includes only heavy atoms from the H2-H5 bundle while bottom figures also include the CTE. (A) RMSD with respect to the first frame of the trajectory. (B) Short range interaction energy between Vt and A3-A5 subunits. (C) BSA between Vt and A3-A5 subunits

**Fig. S2.**
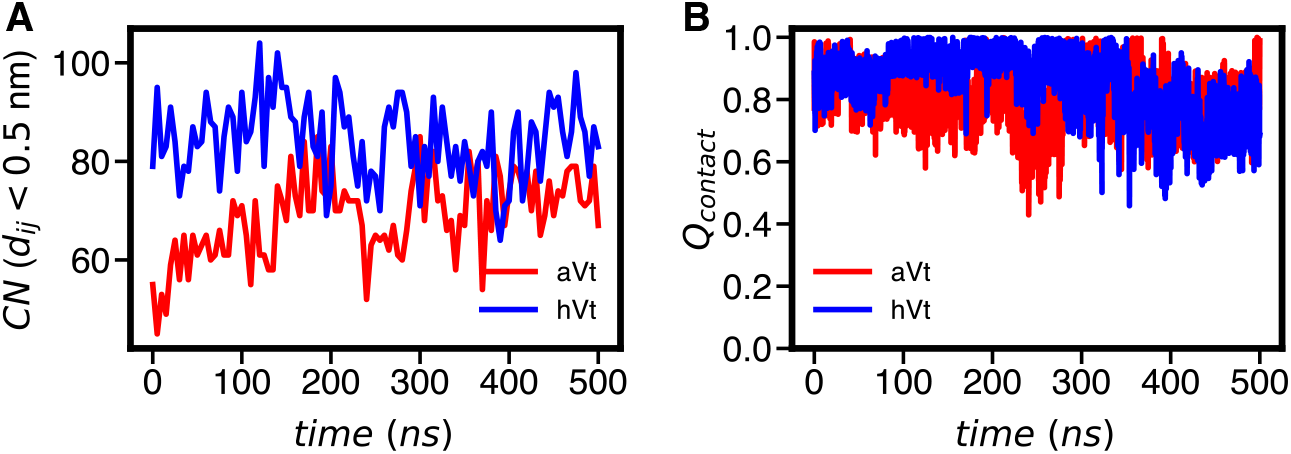
(A) The total number of contacts (CN) between A3A5 and Vt+CTE vs time. (B) *Q*_contact_ vs. time which is only based on distances from H4-H5 residues to the A3/A5 interface.

**Fig. S3.**
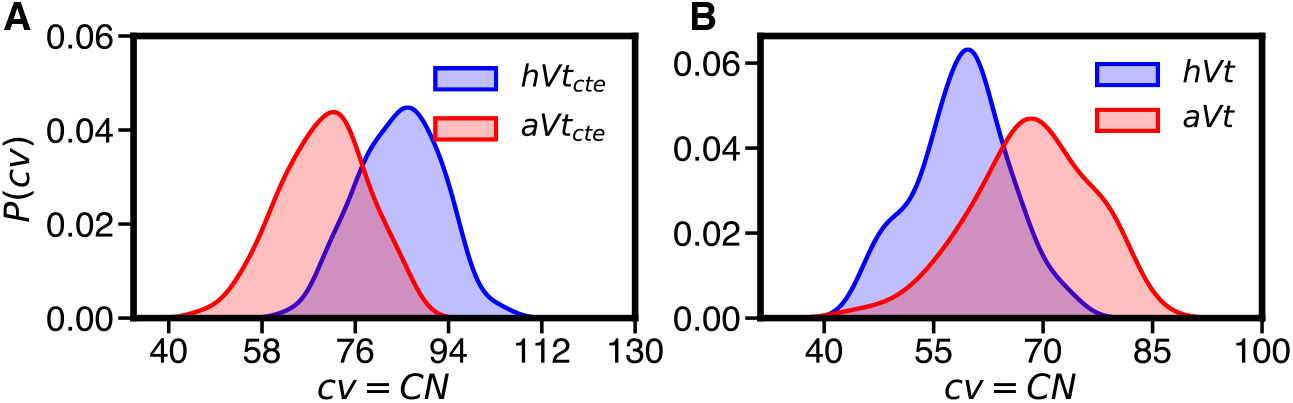
(A) Distribution of the total number contacts between H2-H5+CTE and the A3/A5 interface. (B) Distribution of CN without CTE.

**Fig. S4.**
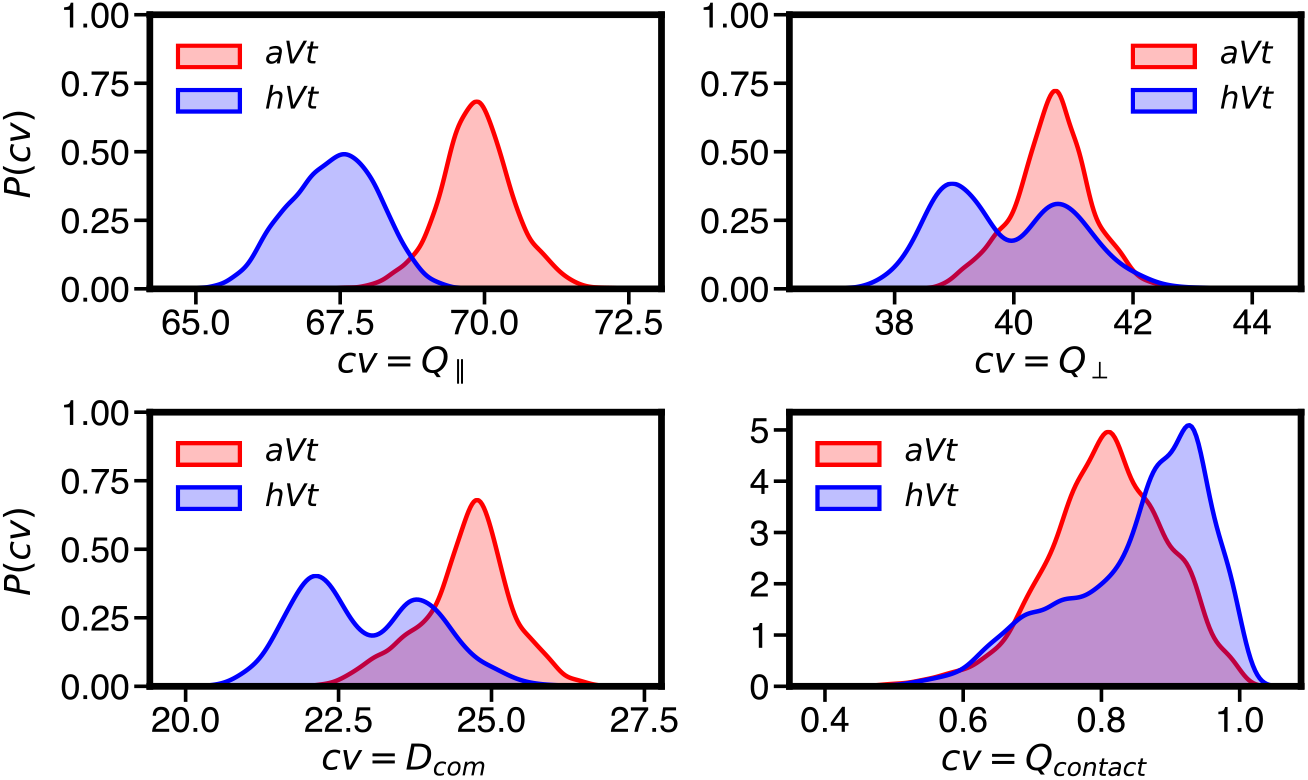
Distribution of CV’s including *D*_com_. which is the distance from the COM of Vt to COM of A3A5 A5-TYR-143-N GLN-1062-OE1 GLN-994-OE1

**Fig. S5.**
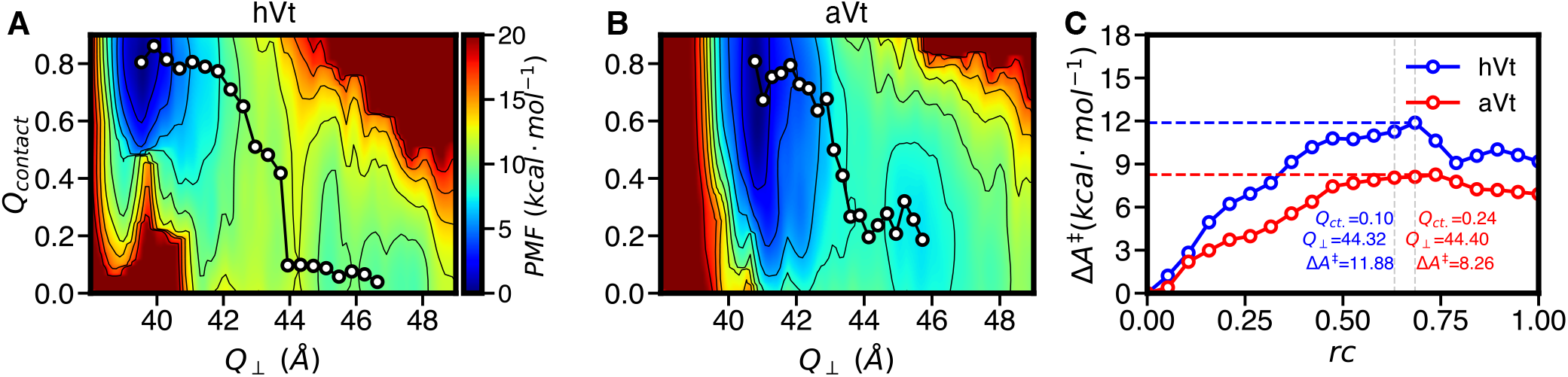
(A,B) FES for the Holo and Aligned models as a function of *Q*_contact_ and *Q*_⊥_ (C) One dimensional projections of the minimum free energy paths from A and B show that Holo is more stable and has a higher barrier to unbinding relative to Aligned. As in Fig. 2 The *Q*_⊥_ and *Q*_contact_ values at the putative transition state are labeled, which are used subsequently to help define the excluded region in unbinding rate calculations.

**Fig. S6.**
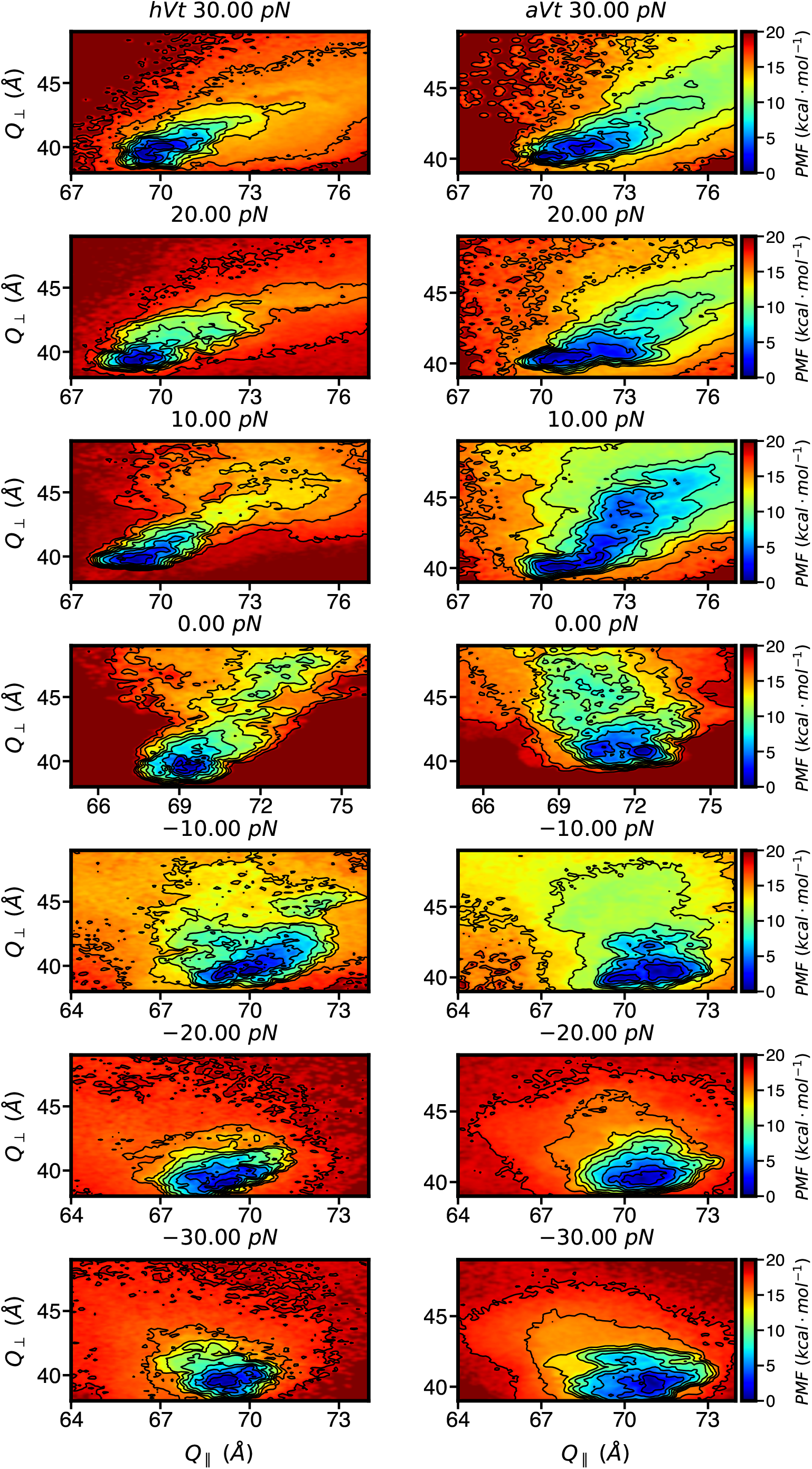
2D FESs for Vt unbinding as forces in either direction for both Holo (left) and Aligned (right) models

**Fig. S7.**
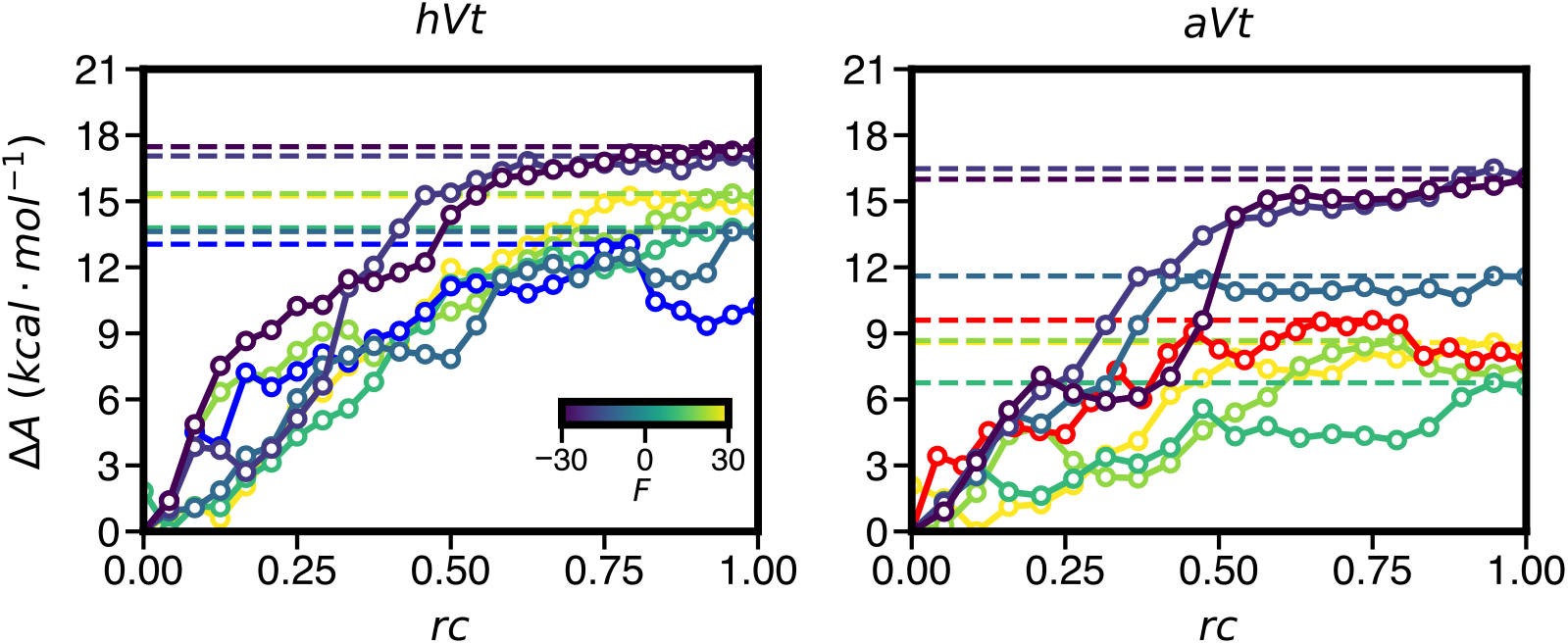
Minimum free energy paths for Vt unbinding at different forces in either direction computed by the string method on the 2D FESs in Fig. S6.

**Fig. S8.**
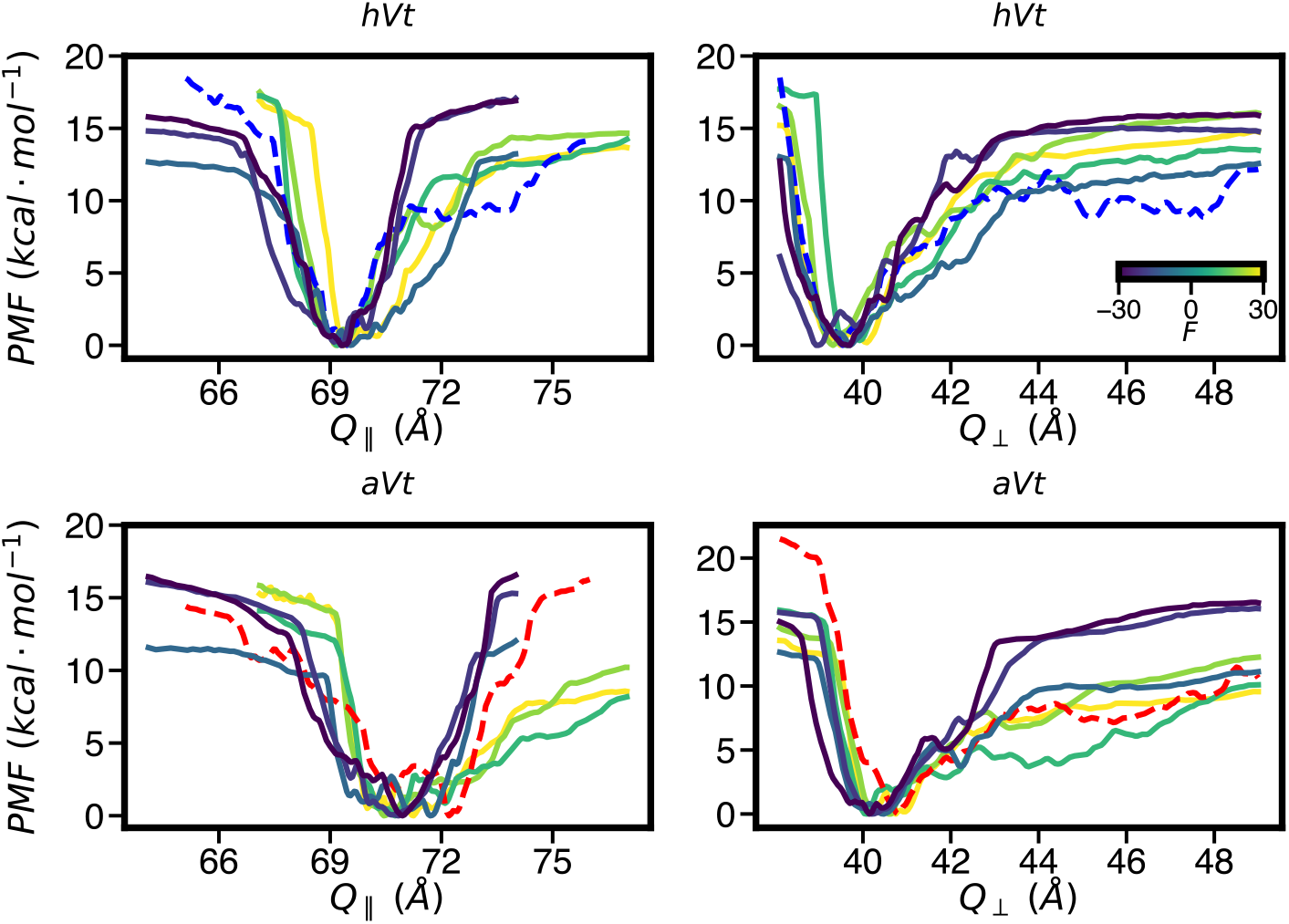
One dimensional projections of the 2D FESs in Fig. S6 at different forces onto individual CVs. Forces are indicated by the color bar, with the dashed line showing *F* = 0.

**Fig. S9.**
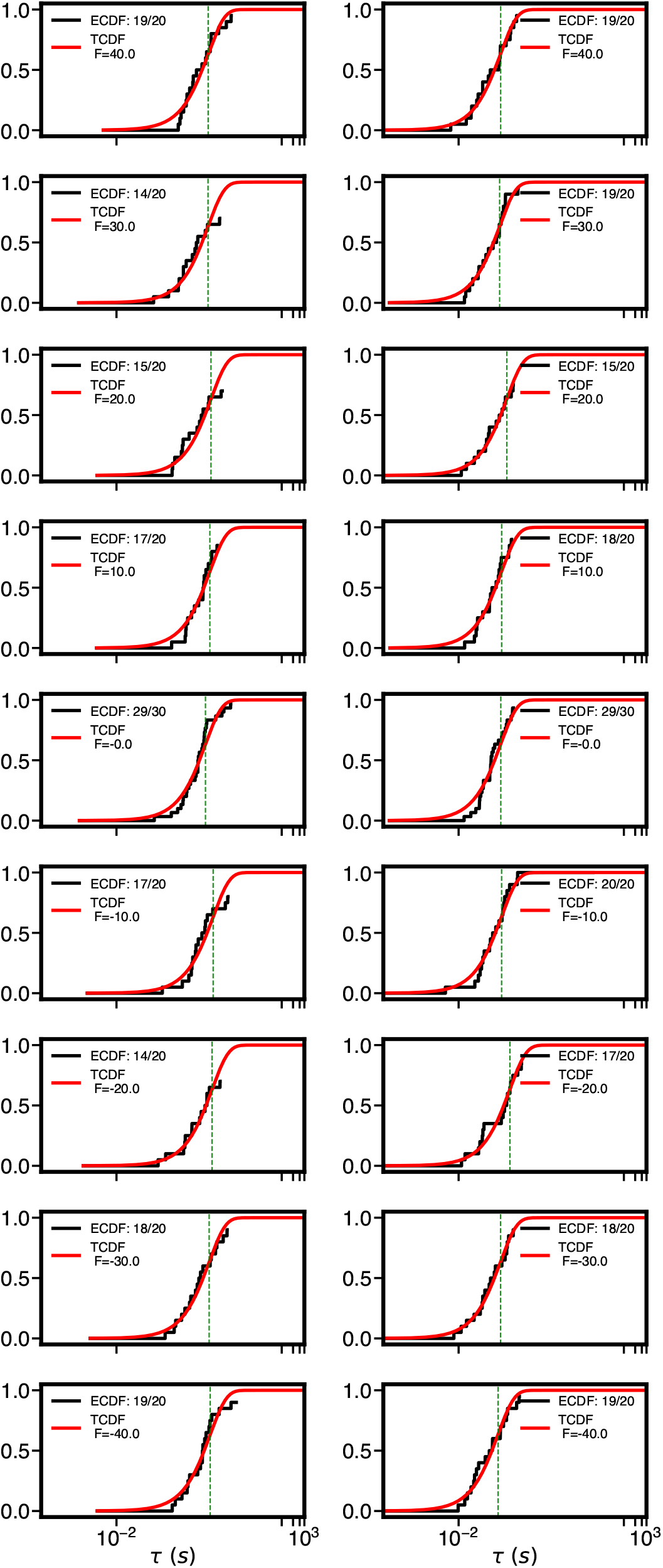
Cumulative distribution functions of rescaled times from OPES-Flooding simulations, corresponding to the lifetimes in Fig. 4. TCDF is the best fit to the data. All fits pass the KS test with a *p*-value > 0.05 for aVt while all but 2 pass the KS test for hVt. The number of runs that reached an unbound state and the total number of runs performed for each condition is given in the caption.

**Fig. S10.**
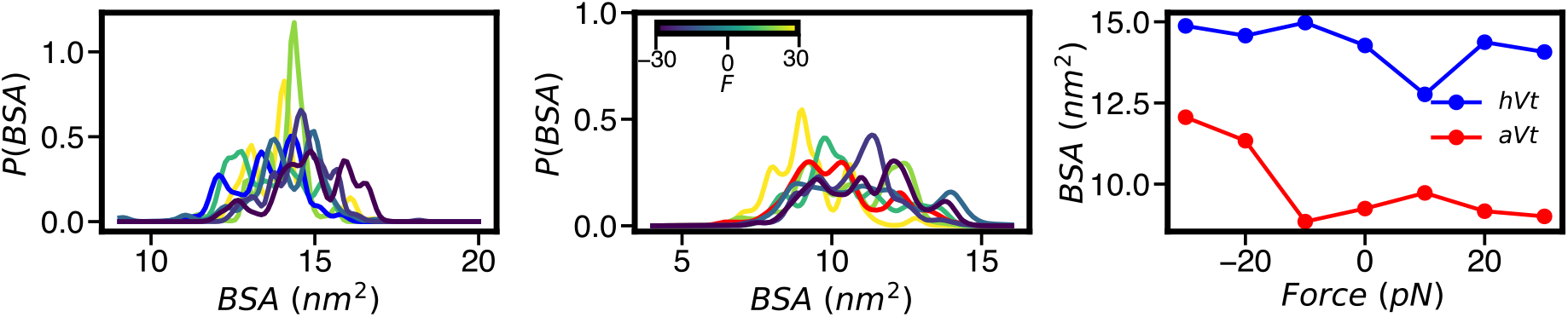
(A,B) Weighted probability distribution of buried surface areas (BSA) for aVt and hVt obtained from FES simulations, BSA’s are computed from SASA’s of heavy atoms of aVt and hVt (including CTE tail) and heavy atoms of actins 3-5 using *BSA* = 0.5(*SASA*_*a*_ + *SASA*_*b*_ − *SASA*_*ab*_), where *a, b*, and *ab* denote the SASA of A3-A5, Vt, and A3-A5 Vt respectively. SASA’s were obtained with GROMACS’ sasa program. (C) Peak BSA’s at each pulling force for aVt and hVt. The BSA’s for aVt are lower than for hVt for all scenarios and BSA’s increase when Vt moves towards the barbed end of F-Actin and remain the same or decrease when Vt moves towards the pointed end.

**Fig. S11.**
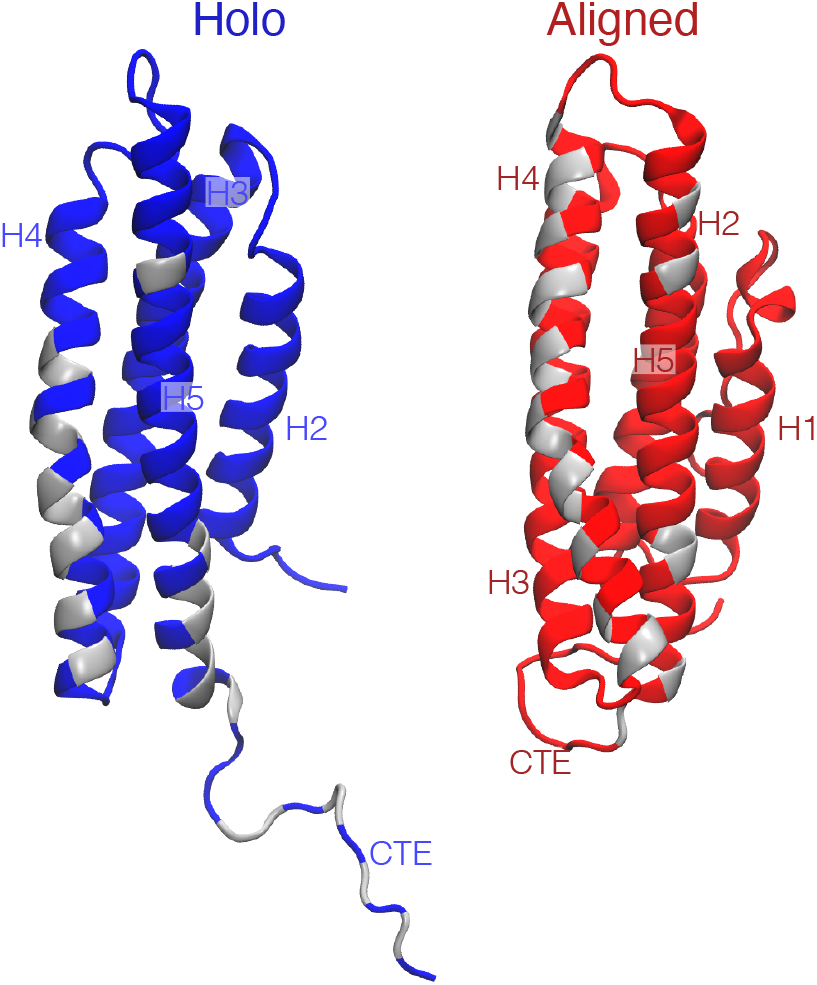
Representative Holo and Aligned stuctures from Fig. 6. Here, atoms in contact with actin are highlighted in silver. The Aligned model has a similar amount of contacts and burried surface area on helices H2-H5 with actin, but they are primarily localized to H4.

## Notes

https://github.com/hocky-research-group/Pena-Vinculin-Unbinding

### Competing Interest Statement

The authors have declared no competing interest.

